# Beneficial and detrimental genes in the cellular response to replication arrest

**DOI:** 10.1101/2021.05.05.442863

**Authors:** Luciane Schons-Fonseca, Milena D. Lazova, Janet L. Smith, Alan D. Grossman

## Abstract

DNA replication is essential for all living organisms, and a variety of events can perturb replication, including DNA damage (e.g., pyrimidine dimers, crosslinking) and replication arrest due to “roadblocks” such as DNA-binding proteins or transcription. Bacteria have several well-characterized mechanisms for repairing damaged DNA and restoring functional replication forks. However, little is known about the repair of stalled or arrested replication forks in the absence of DNA lesions. Using a library of random transposon insertions in *Bacillus subtilis*, we identified 35 genes that affect the ability of cells to survive arrest of replication elongation, in the absence of DNA damage. Genes identified included those involved in iron-sulfur homeostasis, cell envelope biogenesis, and DNA repair and recombination. In *B. subtilis*, and many bacteria, two nucleases (AddAB and RecJ) are involved in early steps in repairing replication forks arrested by DNA damage. Loss of either one causes increased sensitivity to DNA damage. These single-strand nucleases resect DNA ends, leading to assembly of the recombinase RecA onto the single stranded DNA. Notably, we found that disruption of *recJ* increased survival of cells following replication arrest, indicating that, in the absence of DNA damage, RecJ is detrimental to surviving replication arrest. In contrast, and as expected, disruption of *addA* decreased survival of cells following replication arrest, indicating that AddA promotes survival. The different phenotypes of *addA* and *recJ* mutants appeared to be due to differences in assembly of RecA onto DNA. RecJ promoted too much assembly of RecA, and loss of RecA compensated for the detrimental effects of RecJ. Our results indicate that in the absence of DNA lesions, RecA is dispensable for cells to survive replication arrest and the stable RecA nucleofilaments favored by the RecJ pathway may lead to cell death by preventing proper processing of the arrested replication fork.

## Introduction

DNA replication is essential for all living organisms, and a variety of events can perturb replication, including DNA damage and replication arrest due to “roadblocks” such as DNA-binding proteins or transcription. In any of these cases, some or all the components of the replication machinery (the replisome) may disassemble from the complex, leading to the collapse of the replication fork (Cox *et al*., 2000; Michel *et al*., 2007). If a subsequent replisome reaches unrepaired lesions, nicks, or collapsed forks, it can turn these defects into double-strand breaks, which, if not repaired, are lethal.

Proper cell growth and division following replication arrest depends on restart of replication. A wide range of bacteria use a similar mechanism for replication restart, where the DNA is processed to enable the formation of a structure resembling a replication fork (reviewed in Singha et al., 2020). This processing typically involves homologous recombination mediated by RecA (Friedberg *et al*., 2006). To initiate homologous recombination, helicases and nucleases process DNA at the arrested fork to generate single-stranded DNA (ssDNA), in a process called end-resection (Lovett and Kolodner, 1989; Chédin *et al*., 2000).

*Bacillus subtilis* has two nucleases capable of performing end-resection, RecJ and AddAB. In response to DNA damage, the exonuclease activity of RecJ can expand ssDNA gaps by several kilobases (Courcelle and Hanawalt, 1999; Lovett and Kolodner, 1989; Dianov *et al*., 1994). In contrast, the helicase-nuclease complex AddAB, a functional homolog of RecBC in *Escherichia coli*, binds blunt or near blunt DNA ends. AddAB degrades both strands until reaching a Chi site (Chédin *et al*., 1998) where it switches to to degrade only a single DNA strand in the 5’ to 3’ direction (Chédin *et al*., 2000). Both the RecJ and AddAB end-resection pathways in *B. subtilis* lead to loading of RecA onto the DNA by the ‘loader proteins’ RecO and RecR (Lenhart *et al*., 2014). RecA forms nucleofilaments that can search and invade a homologous double-stranded DNA, creating cross-shaped DNA structures known as Holliday junctions. Finally, specific resolvases process Holliday junctions into a substrate for the replication restart protein PriA.

Almost everything known about the bacterial response to replication arrest comes from experiments using DNA damage to disrupt DNA replication. DNA repair and replication restart are both required for optimal survival, making it difficult to differentiate between processes related to the damage itself or the replication fork arrest. We were interested in finding genes that influence the ability of cells to survive replication arrest, without the confounding effects of chemical damage (e.g., pyrimidine dimers, cross-links) to DNA.

To cause replication fork arrest independent of DNA damage, we used HPUra (6-(p-hydroxyphenylazo)-uracil), an azopyrimidine that reversibly binds to and inhibits the catalytic subunit of DNA polymerase III (PolC) in *B. subtilis* (Brown, 1970). We used Tn-seq (transposon insertion mutagenesis with massively parallel sequencing) to identify candidate genes that affect cell survival following the arrest of replication elongation caused by treatment with HPUra.

We found that many processes appeared to be important for survival, including: the regulation of oxidative homeostasis, cell wall stability, and DNA repair pathways. Interestingly, we found that one of the DNA end resection pathways that is important for surviving DNA damage was detrimental to surviving replication arrest. Loss of *recJ, recO*, and *recF* was beneficial to cell survival, indicating that the functional genes were detrimental. In contrast, functional *addA, recN*, and *recU* were beneficial for cell survival following replication fork arrest. Our findings indicate that although AddAB and RecJ both carry out end-resection, they commit replication fork repair to distinct pathways. We found that these pathways led to different amounts of RecA assembled onto DNA. Following replication fork arrest, RecJ or the recombinase loaders led to excessive RecA loading, accumulation of unrepaired forks, and ultimately an increase in cell death. Our data indicate that *B. subtilis* activates at least two pathways to repair arrested replication forks due to replisome malfunction and that one pathway (AddAB) is beneficial and the other (RecJ) is detrimental to cell survival.

## Results

### Identification of genes involved in cell survival following replication arrest

Using Tn-seq, we identified non-essential genes that significantly affected the ability of cells to survive and recover from replication arrest caused by treatment with HPUra. We used a previously described library of ~1.7 × 10^5^ unique transposon insertions of a modified version of the *magellan6* transposon, *magellan6x*, distributed throughout the *B. subtilis* genome (Johnson and Grossman, 2014). This library of cells was split in two and treated or not with HPUra for one hour. The cells were then removed from HPUra, allowing replication and cell growth to resume. Aliquots of cells were harvested 2, 3 and 4 hours later, and the location and relative number of transposon insertions in a given chromosomal site were determined by high-throughput sequencing. In a population of random transposon insertions, mutations that lead to decreased survival would be underrepresented, and the ratio between the frequency of insertions in treated and untreated samples would be <1. Conversely, insertion mutations that improved survival would be overrepresented, resulting in frequency ratios >1.

Thirty-five genes were classified as affecting survival and recovery from HPUra (see Material and Methods for analysis and criteria), and 13 of these were validated using targeted gene disruptions (Table 1). Several cellular processes appeared to affect survival, including: (*i*) iron-sulfur and oxidative homeostasis (*rex, nadABC, defB*, and *nifS*), in agreement with the induction of oxidative stress and genes regulating NAD/NADH following treatment with HPUra (Goranov *et al*., 2005); (*ii*) cell wall stability (*cwlO* and *ponA*); (*iii*) lysogenic phage induction (*yoyI, yonH* from the SPβ prophage and *xpf, xkdBC* from PBSX); and (*iv*) DNA repair and recombination. Below, we focus on several of the DNA recombination and repair genes (Fig. 1A). We also validated seven of the other genes identified in the screen using targeted gene disruptions (Table 1), but did not further evaluate them as part of this study.

**Table 1.**
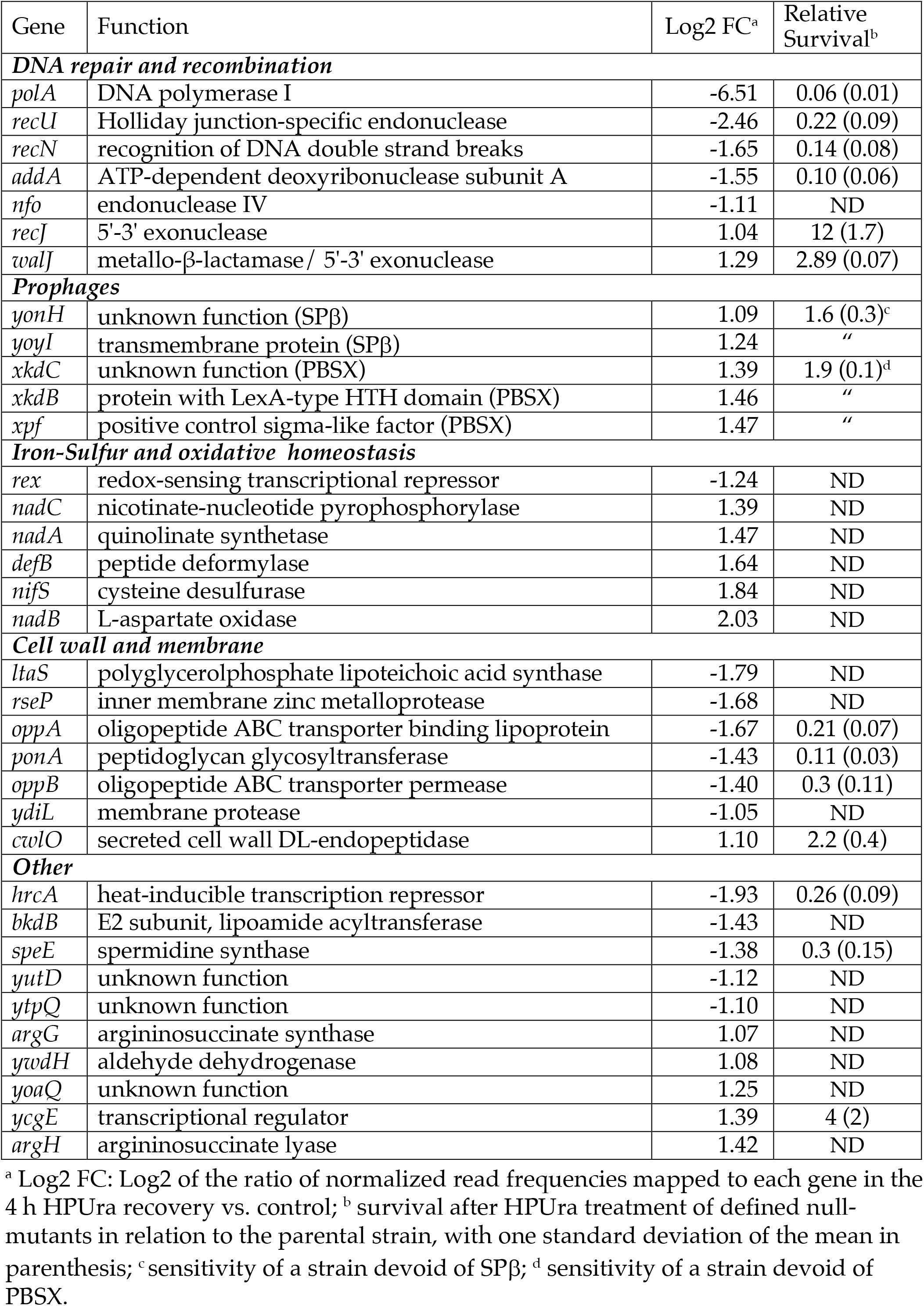
Genes identified in Tn-seq analysis of cells surviving replication arrest in the absence of DNA damage.

**Figure 1.**
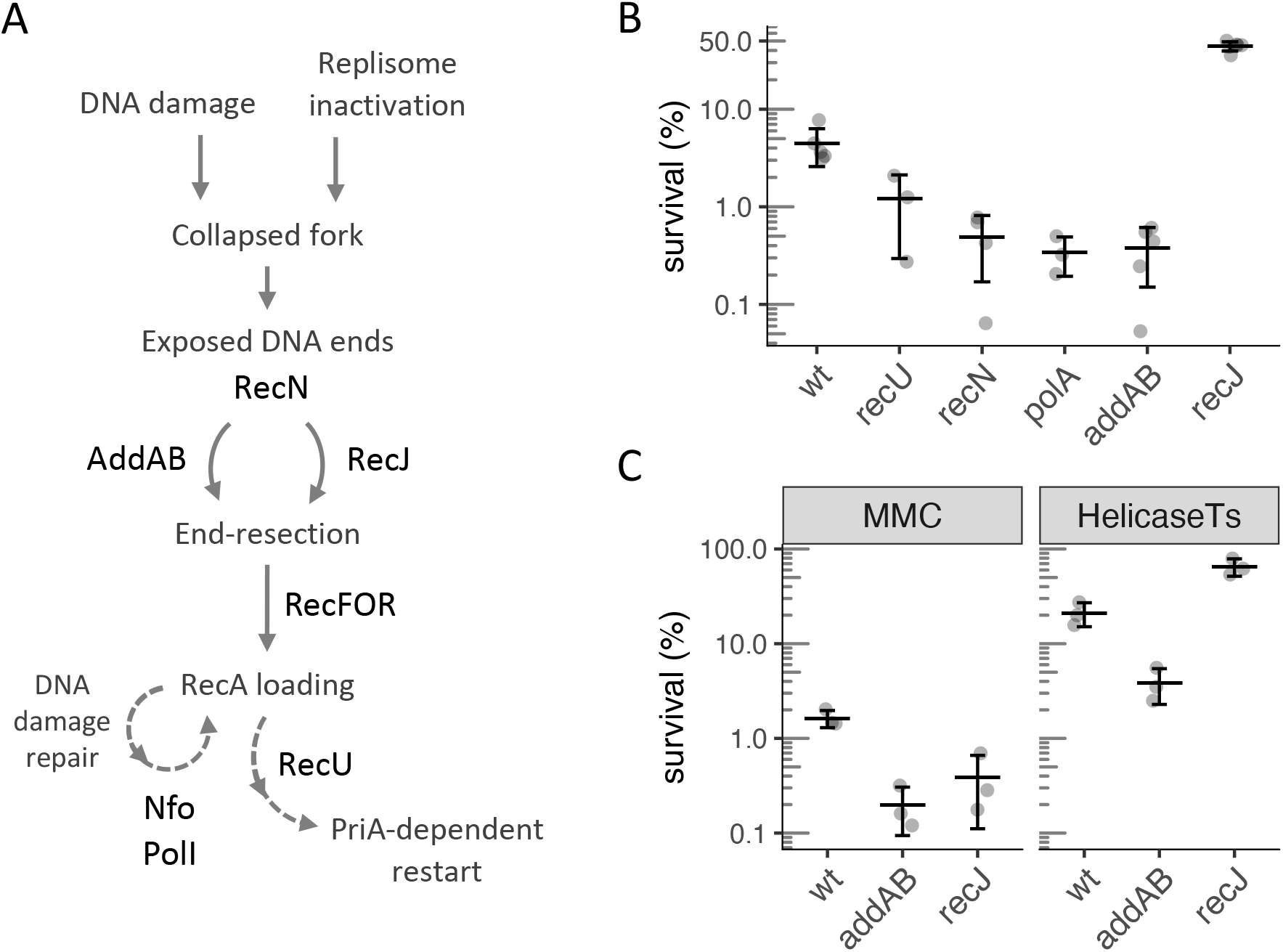
DNA recombination genes affecting survival of DNA damage-independent replication arrest. **A**. General view of the roles of the DNA repair and recombination genes identified in the Tn-seq screen. The dashed arrows represent multistep pathways. **B**. Fold change survival of wild-type (*B. subtilis* strain JMA222) and deletions of *recU* (LSF625), *polA* (LSF041), *recN* (LSF298), *addAB* (LSF253) and *recJ* (LSF200), after 3 h of HPUra treatment, relative to the start of the arrest. All mutants were statistically different from wild-type, as assessed by two-sample t-tests (*P* < 0.025). **C**. Deletion of *recJ* has different effects on survival depending on the nature of replication arrest. First panel: survival of wild-type (JMA222), Δ*addAB* (LSF253) and Δ*recJ* (LSF200) after 3 h of 0.33 µg/ml MMC treatment. Second panel: wild-type, Δ*recJ* and Δ*addAB* cells carrying a *dnaCts* allele (LSF176, LSF274, and LSF326) grown in permissive temperature (30°C) to early exponential phase were shifted to non-permissive temperature (49°C) for 4 h. Fold change in survival is relative to the population before the shift. The deletion mutants in both conditions were statistically different from wild-type, as assessed by two-sample t-tests (*P* < 0.025). Throughout this work, error bars correspond to one standard deviation of the mean and each point is a biological replicate.

In order to confirm the importance of the DNA repair and recombination genes identified in the Tn-seq screen, we made null mutations in six of the loci identified (*polA, recU, recN, recJ, walJ*, and *addAB* (note that *addB* and *addA* comprise an operon and for all experiments except those in Fig. 5 both genes were deleted) and tested their effects on survival after HPUra-mediated replication fork arrest. We treated wild-type and mutant cells with HPUra for 3 h and then removed HPUra and measured the number of viable cells remaining. Approximately 4.5% of wild type cells survived this prolonged replication arrest (Fig. 1B). In contrast, only 0.5-1% of *polA, recU, recN*, and *addAB* mutant cells survived.

In contrast to the detrimental effects of loss of *polA, recU, recN*, and *addAB* on surviving replication fork arrest, we found that loss of *recJ* actually promoted survival. In the *recJ* null mutant, approximately 45% of cells remained viable after 3 hr of replication arrest (Fig. 1B). These results indicate that normally RecJ is detrimental to survival of replication arrest in the absence of DNA lesions. Together, these results confirm the initial Tn-seq data that indicated that these genes are important for promoting survival following replication fork arrest in the absence of DNA damage.

### Comparison of *recJ* and *addAB* mutants in response to replication arrest

Both RecJ and AddAB are involved in end-resection (Fig. 1A), and both improve survival after treatment with DNA-damaging agents (Sanchez *et al*., 2005; Sanchez *et al*., 2006; Carrasco *et al*., 2015). Therefore, it was somewhat surprising that a *recJ*-null mutant had increased survival in response to replication fork arrest.

We confirmed that Δ*recJ* and Δ*addAB* mutants were more sensitive than wild-type cells to treatment with the DNA damaging agent mitomycin C (MMC). Approximately 1.5% of wild-type cells survived treatment with MMC (0.33 µg/ml) for 3 h (Fig. 1C). In contrast, *addAB* and *recJ* mutants had 7 and 4-fold lower survival, respectively.

To verify that the effects of *recJ* and *addAB* on survival and recovery from HPUra were due to replication arrest, rather than some other possible effect, we tested their survival under conditions in which replication was arrested with a temperature-sensitive allele of the replicative DNA helicase (*dnaCts*) (Karamata and Gross, 1970). Inactivation of the helicase mirrored the phenotypes observed during HPUra-mediated replication arrest (Fig. 1C). We grew *dnaCts* (LSF176), *dnaCts* Δ*addAB* (LSF326), and *dnaCts* Δ*recJ* (LSF274) strains at a permissive temperature (30°C). Cells were then shifted to non-permissive temperature (49°C) to arrest replication elongation. After 4 h at 49°C, we measured the number of viable cells from each of the three strains. In the otherwise wild-type cells, prolonged replication arrest at the non-permissive temperature caused a decrease in cell viability to about 20% of the original population (Fig. 1C). The loss of *addAB* exacerbated this effect, and only 4% of cells were viable. In contrast, the loss of *recJ* largely protected the cells from the outcome of replication arrest due to inactivation of the replicative helicase, and 65% of cells survived (Fig. 1C). Together, our results indicate that *recJ* is detrimental to surviving replication fork arrest in the absence of DNA lesions, and confirm previous work (Sanchez *et al*., 2006) demonstrating that *recJ* contributes to survival following DNA damage.

### Deletion of RecA loaders RecO or RecF protects *ΔaddAB* cells from arrest-induced death

End-resection by AddAB or RecJ produces the substrate for interaction with RecA. RecO is the main protein responsible for loading RecA onto the ssDNA substrate (Fig. 1A), and RecF stabilizes the resulting RecA-nucleofilaments (Lenhart *et al*., 2014). Insertions in *recO* and *recF* were not recovered in our initial Tn-seq screen, likely due to harmful effects of natural light on these mutants, causing depletion of insertion mutants in the starting library. Instead, we constructed Δ*recO* and Δ*recF* mutants and tested their effects on survival following replication arrest induced by HPUra treatment.

We found that deletion of either *recO* or *recF* significantly increased cell survival following replication fork arrest (HPUra treatment) in comparison to wild-type (Fig. 2A). Both deletions suppressed the sensitivity of Δ*addAB* mutants to HPUra (Fig. 2A, see *addAB recO* and *addAB recF*), indicating that loading RecA was at least partially responsible for decreased survival of the Δ*addAB* mutant.

**Figure 2.**
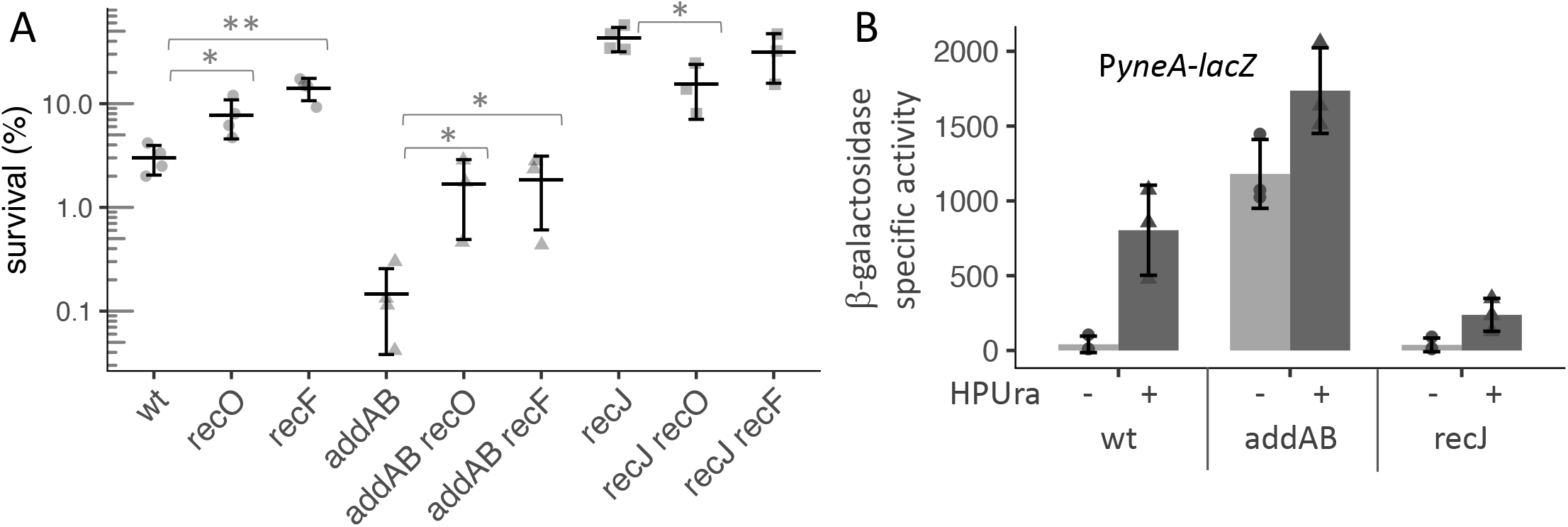
RecA loading is detrimental to survival during HPUra-induced replication arrest. **A**. Survival after HPUra treatment is higher in the absence of RecA loaders. Fold change in survival after 3 h of arrest for wild-type (JMA222), Δ*recO* (LSF814), Δ*recF* (LSF270), Δ*addAB* (LSF253), Δ*addAB* Δ*recO* (LSF817), Δ*addAB* Δ*recF* (LSF709), Δ*recJ* (LSF200), Δ*recJ* Δ*recO* (LSF815), Δ*recJ* Δ*recF* (LSF282). Significant difference in means (two-sample T-test): * *P* < 0.05; ** *P* < 0.01. **B**. RecA activity is lower in the absence of *recJ*. Promoter activation of the SOS-controlled gene *yneA* as a proxy for RecA activity. Cultures of wild-type, Δ*recJ*, and Δ*addAB* strains containing *amyE::*P*yneA-lacZ* (LSF633, LSF634, and LSF635, respectively) were treated or not with HPUra. After 30min, aliquots were collected for measuring β-galactosidase activity. Values are normalized by OD_600_ and represent three independent experiments.

In contrast to effects on the *addAB* mutant, deletion of *recF* had little if any effect on the sensitivity of the Δ*recJ* mutant to HPUra, indicating that RecF and RecJ likely affect the same process – formation of stable RecA filaments. Loss of *recO* appeared to be epistatic to loss of *recJ* (Fig. 2A), consistent with the larger effect of *recO* than *recF* on formation of RecA filaments in response to replication fork arrest with HPUra (Lenhart *et al*., 2014).

### The SOS response is reduced in the absence of *recJ* and increased in the absence of *addAB*

We postulated that if the increased survival of Δ*recJ* is a consequence of lower RecA filament stability, then Δ*recJ* might have lower RecA activity. Once assembled onto ssDNA, RecA induces autocleavage of the SOS repressor LexA, allowing the expression of many genes involved in response to genotoxic stress (Little *et al*., 1980; Wojciechowski *et al*., 1991; Au *et al*., 2005). Consequently, expression of genes repressed by LexA can be used as a proxy for the levels of activated RecA. *yneA* encodes an inhibitor of cell division (Kawai *et al*., 2003), and is repressed by LexA and de-repressed in response to DNA damage and replication fork arrest (Goranov *et al*., 2006).

We used a fusion of the promoter region of *yneA* to *lacZ* (P*yneA*-*lacZ*) and measured expression in the mutants in the presence and absence of HPUra. For wild-type cells in defined minimal medium, β-galactosidase activity from the P*yneA*-*lacZ* fusion increased 7-fold after 30 min of arrest (Fig. 2B). In the Δ*recJ* mutant, there was a basal level of activity similar to wild-type, but only a 2-fold increase after HPUra addition, indicating that *recJ* plays an important role in RecA activation. The absence of *addAB* led to very high P*yneA-lacZ* activity during exponential growth (Fig. 2B), indicating that there is an abundance of active RecA in these cells.

### Lysogenic phages are not responsible for the effects of *recJ* and *addAB* on survival after replication arrest

It seemed plausible that the impact of *recJ* and *addAB* on survival following replication stress could be due to different levels of phage induction, as both mutations affect the activity of RecA. The *B. subtilis* strains used in the experiments described above have two resident lysogenic phages, both of which undergo RecA-dependent activation after genotoxic stress: SPβ and the defective phage PBSX (Craig and Roberts, 1980; Phizicky and Roberts, 1980; Goranov *et al*., 2005; Goranov *et al*., 2006). Both encode toxins and autolysins which, when expressed, can cause cell death. We found that insertions in two genes from SPβ (*yoyI, yonH*) and three genes involved in the early steps of PBSX induction (*xpf, xkdBC*) were overrepresented in the screen, indicating that loss of these genes helped cells survive arrest of replication elongation (Table 1).

Although the presence of SPβ and PBSX contributed to cell death following replication arrest, *addAB* and *recJ* also affected cell survival in cells missing both PBSX or SPβ. We measured survival of cells missing either PBSX, SPβ, or both following replication arrest with HPUra. In these experiments, the survival of wild-type cells (containing both lysogenic phages) was 3.5%, and loss of either phage increased survival to approximately 7%, and loss of both phages increased survival to 15-20% (Fig. 3A).

**Figure 3.**
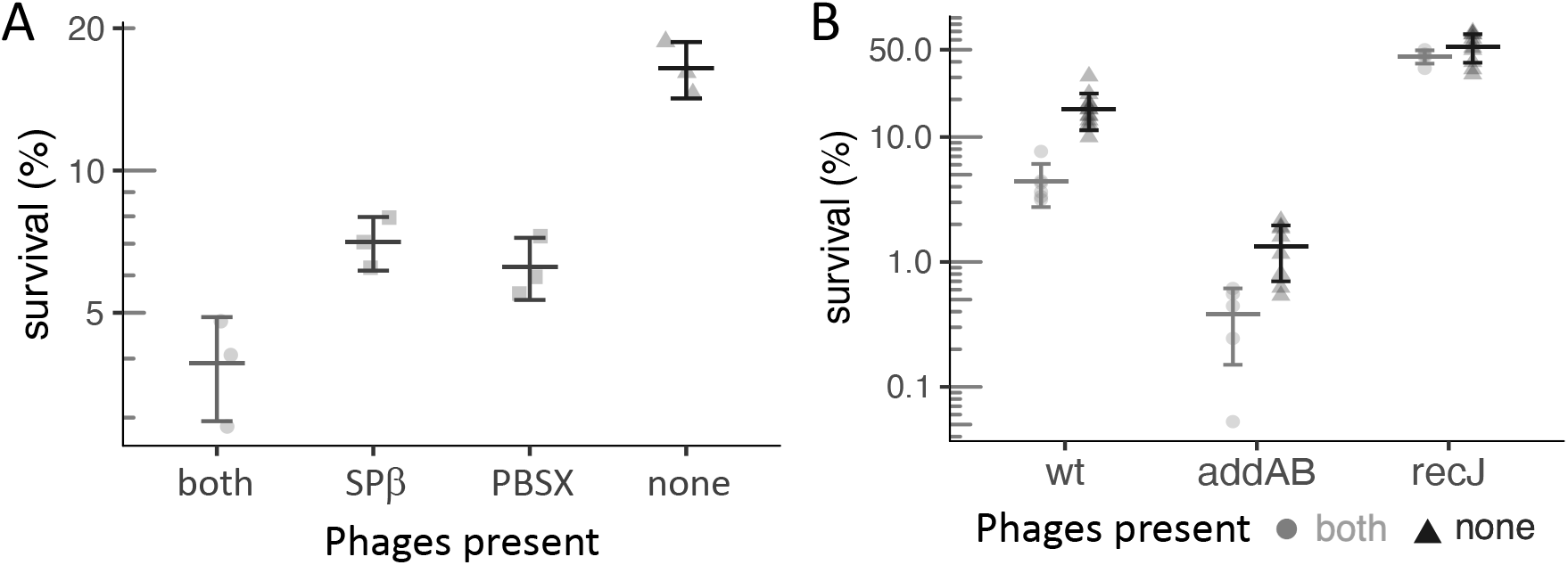
Effect of the resident lysogenic phages SPβ and PBSX on survival after replication arrest. **A**. SPβ and PBSX have an additive effect on cell death after 3 h HPUra treatment. Fold change survival of strains JMA222 (which has both phages), LSF204 (only SPβ), LSF203 (only PBSX), and LSF225 (none). **B**. Δ*recJ* and Δ*addAB* survival phenotypes are independent of phages. Survival of 3 h replication arrest for strains lacking both phage is plotted as dark gray triangles with black error bars; wt (LSF225), Δ*recJ* (LSF231), and Δ*addAB* (LSF254). Data from corresponding strains with both phages (JMA222, LSF253, and LSF200, respectively), plotted as light gray circles with gray error bars, was added for comparison, and are the same as presented in Figure 1B. All strains tested were significantly different from wild-type (two-sample t-test *P* < 0.025).

We found that *addAB* and *recJ* affected survival following replication arrest even in cells missing both PBSX and SPβ. In strains cured of phages, loss of *addAB* caused a 10-fold drop in survival, indicating that this mutant was still more sensitive to replication arrest than wild-type cells (Fig. 3B). This effect was comparable to that caused by loss of *addAB* in strains with both phage (~8-fold). The absence of *recJ* increased the survival of cells after treatment with HPUra to ~50%, regardless of the presence or absence of PBSX and SPβ (Fig. 3B). Together, our results indicate that the effects of *addAB* and *recJ* on survival are largely independent of killing by SPβ or PBSX.

### End-resection promotes survival following replication fork arrest, whereas RecA does not

We sought to determine how strains lacking end-resection or *recA* behaved following replication fork arrest by HPUra. RecA is essential for the survival following treatment with DNA damage agents and, due to the inability to efficiently load RecA in the absence of end-resection, a double Δ*addAB* Δ*recJ* mutant has the same extreme sensitivity to DNA damage agents as Δ*recA* (Lenhart *et al*., 2014; Sanchez *et al*., 2006).

The survival following replication arrest of a strain incapable of performing end-resection (Δ*addAB* and Δ*recJ*), was 0.3% (Fig. 4). This is significantly worse than the effect caused by Δ*addAB* and indicates that the loss of both of the end-resection pathways is synergistic.

**Figure 4.**
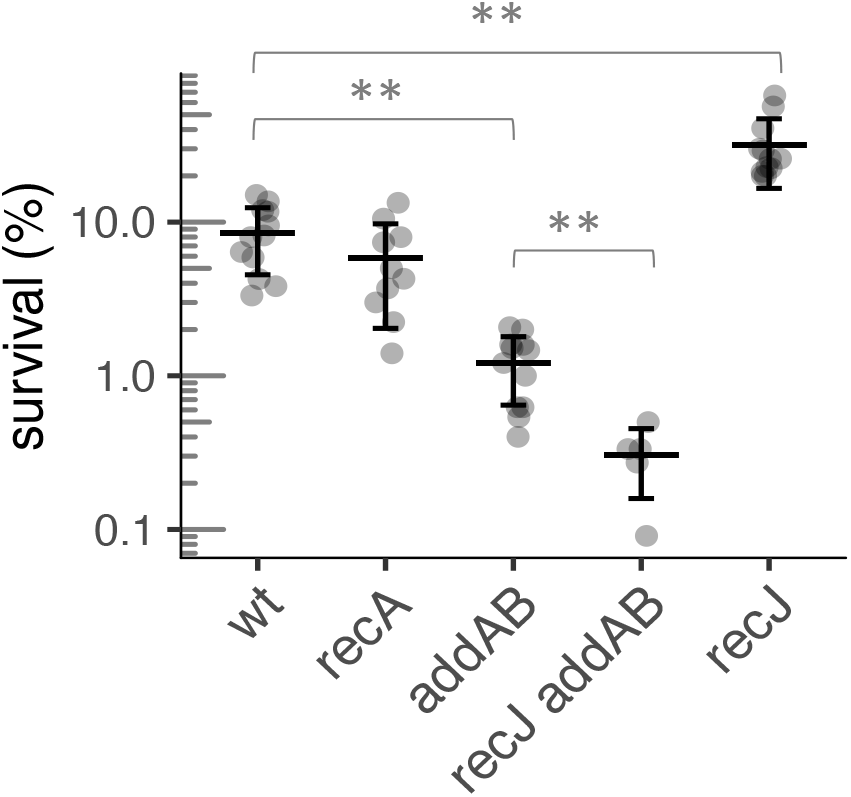
Effect of RecA and end-resection in survival of HPUra-induced replication arrest, in the absence of phages. Fold change in survival after 3 h of arrest for wild-type (LSF225), and strains lacking *recA* (LSF658), *addAB* (LSF254), *recJ addAB* (LSF648), and, *recJ* (LSF231). ** Statistically different means, by two-tailed t-test (*P* < 0.005).

In contrast, *recA* did not detectably affect survival following replication arrest. Survival of the *recA* null mutant was similar to that of otherwise wild-type cells, but lower than the *recJ* mutant (Fig. 4). Based on these results, we conclude that end-resection is essential for recovery from replication fork arrest, but that RecA and RecA-mediated homologous recombination is not required.

### RecA activity correlates with the accumulation of double-strand ends during HPUra-induced replication arrest

Since RecA itself was not required to cope with HPUra-induced arrest, but mutants with lower RecA activity like Δ*recJ* and Δ*recF* had increased survival, we turned to the possibility that RecJ activity and RecA loading was interfering with the proper processing of collapsed replication forks. We reasoned that if that was the case, we would observe the accumulation of recombination intermediates and exposed DNA ends in strains with higher RecA loading. RecN binds to and directs the repair of these ends, coalescing them into a single repair center per nucleoid. When fused to fluorescent proteins, RecN had been used to detect double-strand end formation by fluorescence microscopy (Kidane *et al*., 2004; Sanchez *et al*., 2006).

We constructed strains in which the native *recN* was replaced by *recN-mNeonGreen*, (*recN-mng*) and used them to determine the frequency of repair centers arising from DNA damage-independent arrest. Different mutants containing the fusion were treated with HPUra and scored by the presence of foci after 30 min of replication arrest (Fig. 5A). In wild-type cells growing exponentially, cells with RecN-MNG foci were uncommon (< 2% cells). After 30 min of replication arrest, this frequency reached 20%. In the absence of *recU*, the product of which limits RecA strand exchange (Carrasco *et al*., 2005), or *addA*, this percentage was higher (60% and 40%, respectively). In contrast, only 5-10% of Δ*recJ*, Δ*recF*, or Δ*recO* cells contained foci. A double Δ*recJ* Δ*addA* mutant behaved like Δ*recJ* (Fig. 5A). Finally, deleting *recA* suppressed the induction of double-strand ends by replication arrest (<5% frequency, Fig. 5A). Since RecA is required for their repair (Sanchez *et al*., 2006), the neutral effect of deleting *recA* in survival (Fig. 4) may be a combination of increased survival due to low levels of exposed DNA ends, and cell death due to the inability to repair the breaks or other damages that do occur.

These results support the model that RecJ and the recombinase loaders impede the proper processing of the replication fork via loading of RecA and indicate that the primary source of double-stranded ends following HPUra-induced arrest is the RecJ pathway.

**Figure 5.**
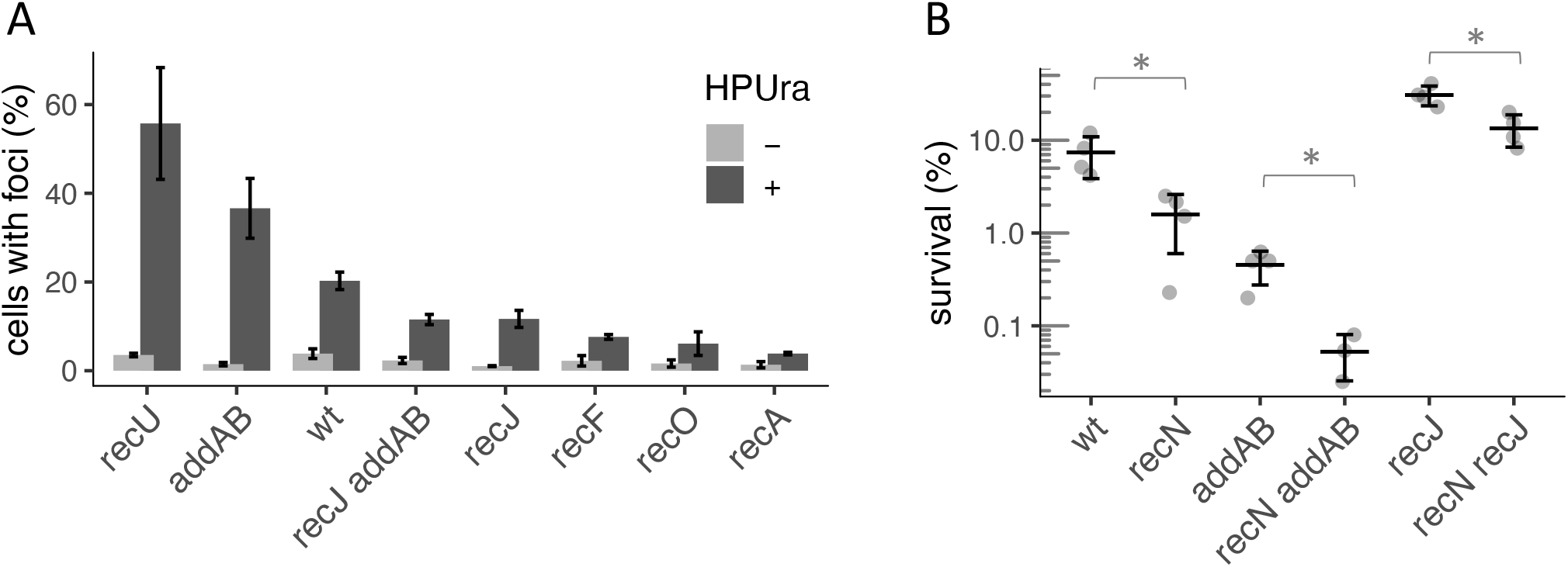
Accumulation of exposed DNA ends during replication arrest by HPUra. **A**. Repair centers in response to exposed DNA ends were visualized by fluorescence microscopy of cells containing RecN-mNeonGreen fusions. Before and after 30 min treatment with HPUra, membranes were stained by FM-464 and nucleoids by DAPI for visualization. At least 800 cells were analyzed for the presence of foci in two biological replicates for wild-type (LSF708), Δ*recU* (LSF740), Δ*addAB* (LSF650), Δ*recO* (LSF818), Δ*recJ* (LSF649), Δ*recJ* Δ*addAB* (LSF706), and Δ*recA* (LSF756). **B**. Additive effects of deleting *addAB* or *recJ* from a Δ*recN* background. Survival of wild-type (LSF225), Δ*recN* (LSF536), Δ*addAB* (LSF254), Δ*recN* Δ*addAB* (LSF652), Δ*recJ* (LSF231), and Δ*recN* Δ*recJ* (LSF651). *Statistically different means according to two-tailed t-test: * *P* < 0.05.

### RecN is important for survival if RecA is assembled onto DNA

Our results indicate that there are dsDNA breaks in wild type cells following replication fork arrest. Furthermore, we found that the loss of *recN* caused a decrease in survival in otherwise wild type cells (Fig. 5B). This decrease in survival was exacerbated by loss of *addAB* (Fig. 5B), consistent with the results indicating that RecA is more active and there are more dsDNA breaks in the absence of *addAB* than in otherwise wild type cells.

In contrast to the phenotype of the *recN addAB* double mutant, the decreased survival of the *recN* single mutant (compared to wild type) was suppressed by loss of *recJ* (Fig. 5B). Again, this is consistent with the notion that RecJ is the major route to loading RecA onto DNA, and that with decreased RecA loading, there is a decrease in dsDNA breaks (Fig. 5A) and a decrease in the need for *recN* in the ability of cells to survive replication arrest.

## Discussion

The replisome pauses or collapses not only when it encounters damaged DNA, but also when roadblocks, such as DNA binding proteins, are encountered, or when the replisome itself is inhibited or damaged. In this paper, we were able to specifically focus on how cells recover from replication arrest by replisome inhibition. The importance of RecA activity and both the RecJ and AddAB end-resection pathways for surviving DNA damage-inducing agents is well established in *B. subtilis* (Courcelle and Hanawalt, 1999; Courcelle and Hanawalt, 2003; Lenhart *et al*., 2012). In contrast, we found a different scenario when there is no DNA damage and a replisome component is inhibited (chemically or using a thermosensitive allele). In this situation, end-resection was necessary, but the RecJ pathway and high RecA activity were detrimental, whereas the AddAB pathway was beneficial. Our results show that different types of genotoxic stresses require different levels of RecA activity, which are obtained by the AddAB vs. RecJ pathways, and that the choice of pathway affects survival.

### The consequences of using each end-resection pathway depend on the kind of replication arrest

As depicted in Fig. 6A, when replication is blocked, both end-resection pathways work together to process the collapsed replication fork. After fork regression, which involves reannealing of the template DNA and annealing of the two daughter strands, AddAB can unwind and degrade the daughter strands until it reaches a *Chi* site, after which it begins generating a 3’ ssDNA tail (Chédin *et al*., 1998). RecJ can target the collapsed fork directly, or an already resected fork (Viswanathan and Lovett, 1998), probably extending the ssDNA substrate generated by AddAB and leading to longer RecA nucleofilaments. RecFOR then ensures RecA nucleofilament loading and stability (Lenhart *et al*., 2014). In this case, the role of RecA is not to promote recombination, but rather to stabilize the DNA at the fork until the DNA damage has been repaired (Courcelle and Hanawalt, 1999; Courcelle and Hanawalt, 2003).

**Figure 6.**
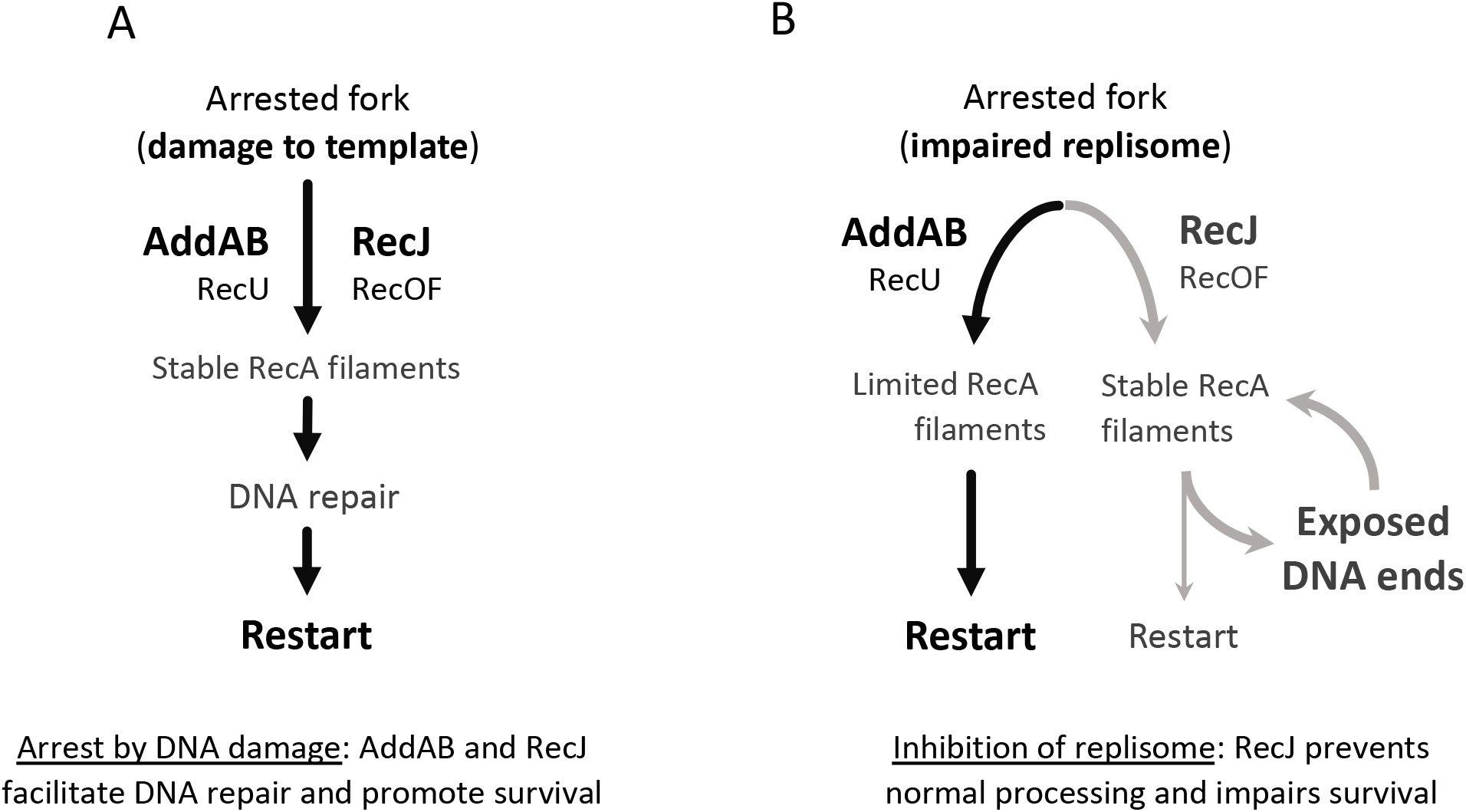
The role of end-resection pathways on survival outcome after replication arrest. **A**. When replication arrest is caused by lesions on the DNA template or other genotoxic stresses requiring RecA activity, both end-resection pathways work together, generating stable RecA filaments that delay restart and support DNA repair. **B**. If replication arrest is caused by the inhibition of replisome components or other genotoxic stresses that do not require RecA, AddAB favors a pathway (black arrows) with limited RecA filaments that promotes survival. RecU helps reducing RecA activity. In this situation, RecJ and the recombinase loaders RecO and RecF impair survival by promoting a pathway (gray arrows) with high RecA activity. This could prevent proper processing of the fork, exposing DNA ends and creating a futile circle.

When replication stops because a replisome component is inhibited, our results indicate that AddAB activity supports optimal recovery (black arrows in the left side of Fig. 6B). After fork regression, AddAB generates less substrate for RecA loading than RecJ. Also, RecU promotes RecA displacement in favor of RuvAB or RecG (Cañas *et al*., 2014), which are capable of faster branch migration (Amit *et al*., 2004). The result of AddAB and RecU activities would be an efficient restart with limited or no RecA participation.

Our data indicate that RecJ and the loaders RecO and RecF negatively affect the fork repair process in this scenario by enhancing RecA activity and preventing fork regression (gray arrows by the right side of Fig. 6B). RecJ can target a range of DNA substrates (Dianov *et al*., 1994; Morimatsu and Kowalczykowski, 2014). In addition to competing with AddAB for access to an already regressed fork, it can degrade a daughter strand in the collapsed fork, which would prevent fork regression and the generation of the blunt ends AddAB requires. Also, RecJ and RecO localize to the replication fork via interaction with SSB (Costes *et al*., 2010), and therefore have immediate access to the replication fork, whereas AddAB does not. In the absence of DNA damage, the resulting stable RecA filaments that help promote survival in other conditions may expose DNA ends that unnecessarily delay fork processing. Any DNA breaks or exposed ends arising during this step would need to go through an additional round of RecN binding, end-resection, and RecA loading, feeding the detrimental pathway.

### Different types of genotoxic stresses require different levels and duration of RecA activity

The different effects that RecA has in survival, depending on the kind of replication arrest, could be due to the complexity of the resulting collapsed fork and duration of RecA activity necessary for restart. Treatment with MMC, for example, impedes replication due to template lesions and “dirty” DNA breaks, *i*.*e*. ones that cannot be ligated because they lack a 3’ hydroxyl or 5’ phosphate. The recovery from this kind of arrest can take up to 3 h in *B. subtilis* and relies on high levels of RecA activity (Kidane *et al*., 2004; Kidane and Graumann, 2005). The repair of a “clean”, ligatable single break in the chromosome, however, requires very transient RecA filaments and takes less than 20 min, at least in *E. coli* (Amarh *et al*., 2018).

When a DNA-binding protein acting as a roadblock arrests replication, *E. coli* cells are able to process these forks and recover normally regardless of the presence of *recA* (Possoz *et al*., 2006). In this case, there are no chemical lesions to be repaired or complex structures to be disassembled at the fork before processing. Likewise, our data indicate that the collapsed forks resulting from the inhibition of a replisome component (PolC by HPUra treatment or DnaC*ts* at the non-permissive temperature) readily undergo fork regression and processing. In this scenario, we showed that the excess RecA activity that results from RecJ and the recombinase loaders is not only unnecessary but also detrimental to survival.

Replication-transcription conflicts also cause replication arrest (reviewed in Merrikh *et al*., 2012, Lang and Merrikh, 2018). The importance of RecA in replication restart at these site is unclear. De Septenville *et al*. (2012) found that RecA was not required in *E. coli*, but Million-Weaver *et al*. (2015) found that RecA was needed in *B. subtilis*. The fact that RecA is required in *B. subtilis* for resolving head-on replication-transcription conflicts, but not for recovery from replication arrest caused by HPUra may be due to the different structures that are formed after each of these treatments. HPUra treatment has been shown to leave many of the replisome components largely intact, at least initially (Liao *et al*., 2015; Liao *et al*., 2016; Li *et al*., 2019), whereas replication-transcription conflicts likely causes replisome disassembly (Mangiameli *et al*., 2017), and this may predispose arrested forks to different restart pathways.

### RecJ leads to exposed DNA ends and increased cell death when fork repair does not require RecA activity

We hypothesize that during HPUra-induced arrest in a wild-type cell, the stable RecA nucleofilaments favored by RecJ prevent optimal repair of the collapsed fork by unnecessarily delaying restart, and by increasing double-stranded breaks. We noted a positive correlation between the predicted levels of RecA activity in different mutants following HPUra treatment and the accumulation of exposed DNA ends. These double-strand ends are probably a combination of unprocessed stalled replication forks and actual breaks. Stable RecJ-dependent RecA filaments may factor into the propensity to cause breaks in these long stretches of coated ssDNA. We found that deletion of *recN* decreased survival of HPUra treatment in wild-type, Δ*addAB*, and Δ*recJ* mutants, suggesting that under all of these conditions double-stranded ends are present and that RecN may be processing them in parallel to end-resection. These two mechanisms – unnecessarily stalled restart, and increased double-stranded breaks – presumably synergistically contribute to cell death during HPUra treatment.

Few studies had focused on the survival outcome of recombination mutants to replisome inhibition. Nevertheless, there are some hints of a disadvantageous role of RecJ in situations where RecA is not required in *E. coli*, and could be explained by excessive loading of the recombinase. A strain lacking *recB* (analogous to Δ*addAB* in *B. subtilis*) can still process a fork after collapse due to a roadblock, but survival was low, while a strain lacking *recA* was indistinguishable from wild-type (Possoz *et al*., 2006). Although a mechanism for these findings was not proposed, the results suggest that a pathway parallel to RecBCD (presumably RecJ/gap repair) may be processing the collapsed forks in a way that decreases viability. Also, RecJ/RecFOR/RecA have been implicated in preventing fork regression in an *E. coli* DNA polymerase III mutant (*dnaE*ts) at non-permissive temperature, negatively affecting survival (Florès *et al*., 2005).

RecJ homologs are widespread from bacteria to eukaryotes (Sanchez-Pulido and Ponting, 2011). It is not known if the detrimental nature of RecJ and excessive loading of RecA in response to replication arrest is widespread, but the results from *E. coli* discussed above, and the conserved nature of RecJ indicate that this might be the case.

### Bacteria use all pathways for repair, even when it hinders proper recovery

Each end-resection mechanism seems adapted to a kind of genotoxic stress in *B. subtilis*: AddAB for double-strand breaks from damage or fork collapse, and RecJ for gaps from DNA repair. However, our data highlighted that bacteria may use an unfavorable (“wrong”) pathway for repair. RecJ commits repair of some DNA damage-independent collapsed forks to a pathway that decreases viability, and it probably competes with AddAB for blunt DNA ends. RecJ and stable RecA loading only promote survival during DNA repair, but our data indicate that they are also used during replication stresses that do not require RecA. Ultimately, this indicates that there the cells do not have an effective mechanism to discriminate between thes different types of broken forks, potentially leading to increased cell death in some, but enabling robust repair and restart in other circumstances.

## Materials and Methods

### Bacterial strains

All *B. subtilis* strains used in this work derive from JMA222 (Auchtung *et al*., 2005), a version of JH642 (Perego *et al*., 1998) cured of the integrative and conjugative element ICE*Bs*1. Strain genotypes and origin of different alleles are listed in Table 2. Previously described alleles were introduced into JMA222 by transforming naturally competent cells with genomic DNA from the strain of interest. After backcrossing, alleles were combined as required.

**Table 2.**
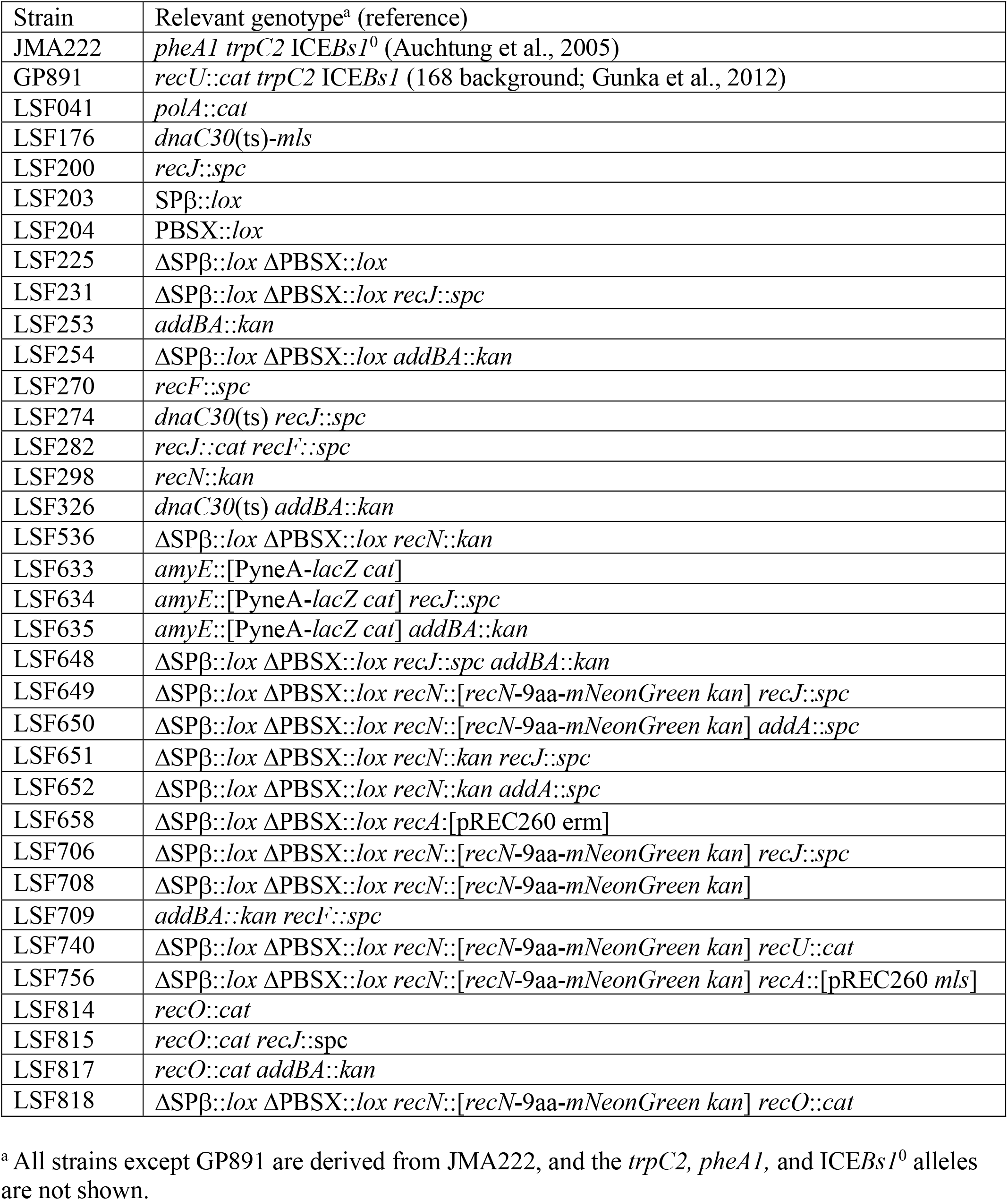
*B. subtilis* strains used.

New strains were constructed by transforming cells with linear DNA products generated by isothermal assembly (Gibson *et al*., 2010). Insertion-deletion constructs contained antibiotic resistance cassettes flanked by 800-1000 bp of genomic sequences upstream and downstream of region to be deleted.

To generate deletions of phages SPβ (LSF203) or PBSX (LSF204), each was substituted by a kanamycin resistance gene flanked by *loxP* sites, generating strains LSF197 (PBSX::*lox-kan-lox*) and LSF198 (ΔSPβ::*lox-kan-lox*). The cassettes were removed by the bacteriophage P1 Cre recombinase using the *cre* expression vector pDR244 as previously described (Meisner *et al*., 2013), leaving a 75 bp scar. A strain devoid of both phages (LSF225) resulted from transforming LSF203 (ΔSPβ::*lox*) with genomic DNA from LSF198 and recombining the antibiotic cassette out.

To visualize exposed DNA ends, we constructed a strain with a *recN*-*mNeonGreen* (*recN-mng*) fusion, inserted in the native locus. The full linear DNA product contained the 800 bp 3’ end of *recN*, an in-frame linker (5’-CTCGAGGGATCTGGCCAAGGAAGCGGC-3’), mNeonGreen coding sequence (a gift from Ethan Garner), a kanamycin resistance cassette, and 800 bp genomic sequence downstream of the *recN* stop codon. Transformation of this construct into JMA222 generated strain LSF284. The *recN*-*mng kan* allele from LSF284 was then introduced into LSF225 (generating LSF708) and null-mutants of interest.

### Media and antibiotics

All experiments were performed in defined minimal medium containing 50 mM MOPS (S7_50_) and supplemented with 1% glucose, 0.1% glutamate, 40 μg/ml phenylalanine and 40 μg/ml tryptophan (Jaacks *et al*., 1989). When required, the following concentration of antibiotics were used: 5 mg/ml chloramphenicol, 100 μg/ml spectinomycin, 100 μg/ml kanamycin, and 0.5μg/ml erthyromycin plus 12.5 μg/ml lincomycin to select for macrolide-lincosamide-streptogramin B (MLS) resistance. Serial dilutions of cultures for spot-plating were made in Spizizen minimal salts medium (Harwood and Cutting, 1991).

### Viability assays

*B. subtilis* strains were streaked from −80°C freezer stocks on LB plates and grown overnight at 37°C. Single colonies were transferred to S7_50_, and dilutions of those were grown overnight with vigorous shaking at 37°C. Starter cultures between OD_600_ 0.05-0.5 were diluted to OD_600_ 0.025, grown for at least three generations, and adjusted to OD_600_ 0.1. All cultures in experiments that included a *recA* deletion were grown in the dark (Δ*recA* leads to light sensitivity due to inability of repairing photo lesions). 38 µg/ml HPUra (6-(p-hydroxyphenylazo)-uracil; Lenhart *et al*., 2014) or 0.33 µg/ml mitomycin C (Sigma-Aldrich) was added, and cultures were kept at 37°C with shaking for 3 h. Viability before and every hour of treatment after was assessed by making 10-fold serial dilutions from 100 μl of cultures in a 96-well plate and plating 10 μl spots on LB in triplicate. Results are in percentage of colony forming units present after 3 h treatment in relation to immediately before addition of MMC or HPUra.

All the growth steps mentioned above were done at 30°C for the viability assays using the temperature sensitive allele of the replicative DNA helicase, *dnaCts* (Karamata and Gross, 1970). At OD_600_ 0.1, the cultures were shifted to 49°C, the non-permissive temperature, to arrest DNA replication. Dilutions of the cultures were plated before and every hour after the shift to assess survival. Results are percentage of colony forming units present after 4 h in relation to immediately before the temperature shift.

Statistical comparison between different groups was performed in R, using the *varequal* and *t*.*test* functions. Two-tailed, paired T-tests were computed using a pooled estimate of variances for similarly distributed samples or were estimated separately for both groups and the Welch modification to the degrees of freedom was used.

### Tn-seq screen

The preparation of the transposon insertion library used in this work is described in detail elsewhere (Johnson and Grossman, 2014). Briefly, MarC transposase was used to catalyze, *in vitro*, the transposition of the *magellan6x* transposon into *B. subtilis* genomic DNA. The transposed genomic DNA was then used to transform JMA222. Transformants were pooled and stored in 30% glycerol.

An aliquot of the transposon insertion library of mutants was diluted in minimal media, and expanded for roughly four generations (doubling time ~ 55min), from OD_600_ 0.02 to 0.3. Half the culture was untreated, while the other had replication arrested by the addition of 38 μg/ml HPUra. After 1 h at 37°C, cells from both cultures were harvested by centrifugation, washed, and resuspended in fresh minimal media, diluting the cultures back to OD_600_ 0.15. They were allowed to recover for 4 h, and aliquots were harvested every hour during this time. DNA was prepared for sequencing as described previously (Johnson and Grossman, 2014), and carried out in an Illumina HiSeq by the MIT BioMicro Center.

### Tn-seq data analysis

The initial processing of Tn-seq data for the removal of sequencing tags from Fastq files was performed according to previous description, using custom-made Python scripts (Johnson and Grossman, 2014). Reads were mapped to JMA222 genome using Bowtie2 (Langmead and Salzberg, 2012). The resulting files contained the number of reads per genomic coordinate in each sample. Our resulting mapped libraries had approximately 10^5^ independent insertions each, with an average of one insertion per 37 base pairs, and 19 insertions per non-essential gene.

Subsequent analyses and file manipulations were performed using custom-made R scripts. Any genomic position with less than 3 reads was discarded, to avoid potential noise due to rare insertions or misaligned reads. Then, we used inter-sample quantile normalization from the preprocessCore package (available at www.bioconductor.org) to ensure that different samples were comparable. Finally, we calculated the number of reads interrupting each gene in every sample – only insertions mapping to 5-95% internal sequence were considered, since insertions in the extremities of essential genes can sometimes be tolerated.

To assess enrichment or depletion of insertion mutants, we calculated the ratio between the number of reads in the treated and control samples, reported as log2 fold change. For proper analysis of Tn-seq, it is important that libraries being compared were expanded roughly the same number of generations before sequencing. Since replication and cell division in the treated library was arrested during 1 h (leaving them one generation “behind” the control libraries), we compared the HPUra-treated libraries with the controls harvested an hour earlier: the HPUra sample harvested after 4 h of recovery was compared to the control harvested after 3 h, and HPUra 2 h with control cells harvested after 1 h of recovery.

Finally, for a gene to be considered as affecting survival in HPUra in relation to the control, it had to satisfy the following requisites: (*i*) be longer than 200bp; (*ii*) have more than 5 insertions in the corresponding control library; (*iii*) have log2 fold change > 1 at 4 h; (*iv*) have an amplified change in read frequency over time. These criteria aimed to restrict the number of candidate genes and decrease false-positives.

### β-Galactosidase activity assay

Cultures in mid-exponential phase growing in minimal media were diluted to OD_600_ 0.1. One-milliliter aliquots were immediately permeabilized with 15 μl toluene and stored at −20°C. HPUra was then added to cultures to 38 μg/ml final concentration. After 30 min at 37°C, 1.5 ml aliquots of the treated cultures were taken, washed by centrifugation at 3000 xg for 2 min, and resuspended in 1.5 ml of fresh media. One milliliter aliquots were permeabilized and frozen, and the remainder was used to measure OD_600_. β-Galactosidase-specific activity was determined similarly to previously described (Miller, 1972). Briefly, 0.5 ml lysates were added to 1 ml ortho-Nitrophenyl-β-galactoside (ONPG) solution. After 1-6 h, reactions were stopped by the addition of 0.5 ml 1M sodium bicarbonate. Cell debris were removed by centrifugation and the supernatants used to measure OD_420_. β-Galactosidase-specific activity was calculated as 1000 × OD_420_ / (min × ml × OD_600_).

### Fluorescence microscopy

To measure exposed DNA ends using RecN-mNeonGreen as a marker, cells containing this construct were grown in minimal media until OD600 ~ 0.1. For each culture, an aliquot was added to a tube containing 4′,6′-diamidino-2-phenylindole (DAPI, 1 μg/ml final concentration) and FM4-64 (2.5 μg/ml), to stain nucleoids and membranes, respectively. After 10 min at 37°C, cells were prepared to be imaged. The remaining of the cultures were treated with 38 μg/ml HPUra for 20 min, when the same dyes were added. Cells were put back at 37°C for 10 min more and then imaged. For imaging, cultures were concentrated roughly four-fold by centrifugation (2 min 2000 *xg*) and 2 μl cells were placed on a slice of 1.5% UltraPure agarose (Invitrogen) dissolved in minimal media. The agarose slice was placed, cells down, on standard coverslips and imaged on a Nikon Ti-E inverted microscope. Fluorescence was generated using excitation/emission of 500/535 nm for RecN-mNeonGreen (1 ms exposure), 350/460 nm for DAPI-stained nucleoids (100 ms) and 510/630 for FM4-64-stained membranes (100 ms). Image processing was performed on Fiji (Schindelin *et al*., 2012), where foci were detected with the Laplacian of Gaussians (LoG) detector in the TrackMate plugin (Tinevez *et al*., 2017), with a 0.7 nm blob diameter and threshold between 10 and 20 (determined by the control samples).

## Acknowledgements

We thank Ethan Garner and Alexandre Bisson-Filho for providing the mNeonGreen allele and initial help with microscopy; Mary Anderson for helpful comments on the manuscript; and Lyle Simmons for providing HPUra. This work was supported by National Institute of General Medical Sciences of the National Institutes of Health under award numbers R37 GM041934 and R35 GM122538 to ADG.

